# Accelerated vaccine induction of IgM neutralising antibody enables vaccine protection against same day lethal influenza virus challenge

**DOI:** 10.1101/2021.02.17.431751

**Authors:** Yoshikazu Honda-Okubo, Lei Li, Jeremy Baldwin, Nikolai Petrovsky

## Abstract

Novel influenza strains to which humans have no pre-existing immunity can trigger global pandemics without warning. Current pandemic vaccines typically require two doses and up to 6 weeks to induce protective immunity. In addition, to their role in increasing vaccine-induced immune response, adjuvants may also play a part in reducing the time between immunization and vaccine protection, although this role has seldom been previously explored in literature. This study assessed the speed of protection achievable with a standard inactivated influenza vaccine when formulated with or without a novel delta-inulin adjuvant (Advax). When formulated with Advax adjuvant mice were protected even when the vaccine was administered intramuscularly contemporaneously with a lethal intranasal virus challenge. The protection was found to be B-cell dependent and transfer of day 6 immune serum from mice immunised with Advax-adjuvanted influenza vaccine conferred protection to naïve animals. This protection was shown to be mediated by vaccine induced IgM rather than IgG neutralising antibodies. The results show that influenza vaccine can be formulated to provide immediate protection following immunization with this novel concept warranting testing in human trials.

**IMPORTANCE:** In the past 100 years there have been several major influenza pandemics that resulted in significant loss of life. The time taken for individuals to develop vaccine protection is an important factor in curbing the spread of infection and reducing pandemic mortality rates. Current influenza vaccines can take up to 5-6 weeks to generate full protection leaving front-line heath care workers and others, at risk for an extended period of time. Our novel accelerated vaccine protection approach provides effectively immediate vaccine protection against lethal virus challenge. This would assist front-line workers to continue to provide essential services and maintain critical infrastructure during pandemic. The study also highlights the often-overlooked role that antigen-specific IgM plays in virus protection and provides a novel adjuvanted strategy for enlisting IgM to provide accelerated vaccine protection.

## INTRODUCTION

Influenza remains a serious public health threat and kills up to 650,000 people worldwide annually according to the World Health Organisation (1). In addition to seasonal epidemic influenza, pandemics caused by novel influenza strains have occurred 5 times in the last hundred years, resulting in a significant loss of life and impacting the global economy (2). The most recent pandemic, A(H1N1)pdm09, in 2009 rapidly spread to over 74 countries within 2 months after its initial appearance in North America (3). Due to the ability of influenza to rapidly develop resistance to antiviral drugs and antibodies (4), vaccination is the single most effective strategy against influenza. However, unlike drug treatment in which the mode of action is immediate, vaccines can take several doses over many weeks to achieve protection. In a pandemic time is a critical factor, highlighting the need to develop vaccine formulations that can reduce the lag time between initial vaccination and immune protection.

Only a limited number of papers in the literature have explored early vaccine protection (0-24hr) against influenza. Mixing of an influenza challenge virus with a live attenuated vaccine reduced disease severity through an unknown mechanism, involving either stimulation of innate immunity and/or interference of the vaccine virus with the challenge virus via genetic reassortment (5). Another study evaluated early protection of an inactivated H5N1 vaccine with an imiquimod adjuvant showed no protection immediately after vaccination and only 35% protection if challenged 1 day after vaccination (5). An inactivated H1N1 vaccine similarly provided only modest protection mediated by innate immunity to a sublethal challenge (6). Hence the literature to date has suggested that it is not possible to sufficiently accelerate adaptive immunity in order to provide vaccine protection against an immediate virus challenge, with only innate immunity being invoked as a source of relatively weak early protection.

Advax is a novel polysaccharide adjuvant based on semi-crystalline microparticles of delta inulin that has a proven track record of boosting vaccine responses and enhancing protection across a wide range of infectious diseases including Influenza (7) (8) (9), Japanese Encephalitis (10), West Nile virus (11), Ebola (12), SARS CoV (13), MERS CoV (14), tuberculosis (15) and anthrax (16), amongst many others. Notably, Advax adjuvanted H5N1 influenza vaccine, after just a single vaccine dose reduced virus shedding and provided robust protection of immunized ferrets against H5N1-associated mortality and clinical disease (9). Advax adjuvant had similar effects in clinical trials of seasonal and pandemic influenza vaccines (17, 18), inducing Day 7 plasmablasts demonstrating high rates of mutations in the B cell receptor hyper-variable region consistent with enhanced B-cell affinity maturation (19). Such increased diversity of the antibody repertoire has been shown to be important to influenza heterosubtypic immunity (20).

Based on the need for more rapidly protective pandemic vaccines, we asked whether Advax adjuvant could accelerate the onset of protective immunity against an influenza vaccine. The current study demonstrates that inactivated influenza vaccine when formulated with Advax was able to provide high levels of protection even if vaccination occurred at the same time as virus infection. Unlike previous studies reported in literature, the accelerated vaccine protection was mediated by the ability of the delta inulin adjuvant to induce an extremely rapid adaptive immune response, with protection mediated by influenza-specific IgM antibodies, rather than innate immunity.

## RESULTS

### Influenza vaccine formulated with Advax adjuvant formulation provides complete protection against lethal challenge 14 days post-immunization

To test the efficacy of a single dose of inactivated iPR8 influenza vaccine with or without Advax adjuvant against lethal challenge with PR8 infection after a standard 14-day post-immunization period, vaccinated BALB/c mice were given 8 x LD_50_ of live virus intranasally 14 days after single-dose vaccination **(Figure 1A)**. Sera collected 14 days after immunization and just prior to virus challenge showed the Advax-adjuvanted iPR8 animals had significantly higher serum influenza-specific IgM and IgG antibodies than animals immunised with iPR8 alone **(Figure 1B)**. Advax-adjuvanted iPR8 immunised mice challenged 2 weeks after immunisation demonstrated complete protection as measured by survival and post-challenge weight loss when compared to iPR8 vaccine alone controls **(Figure 1C-D)**.

**Figure 1.**
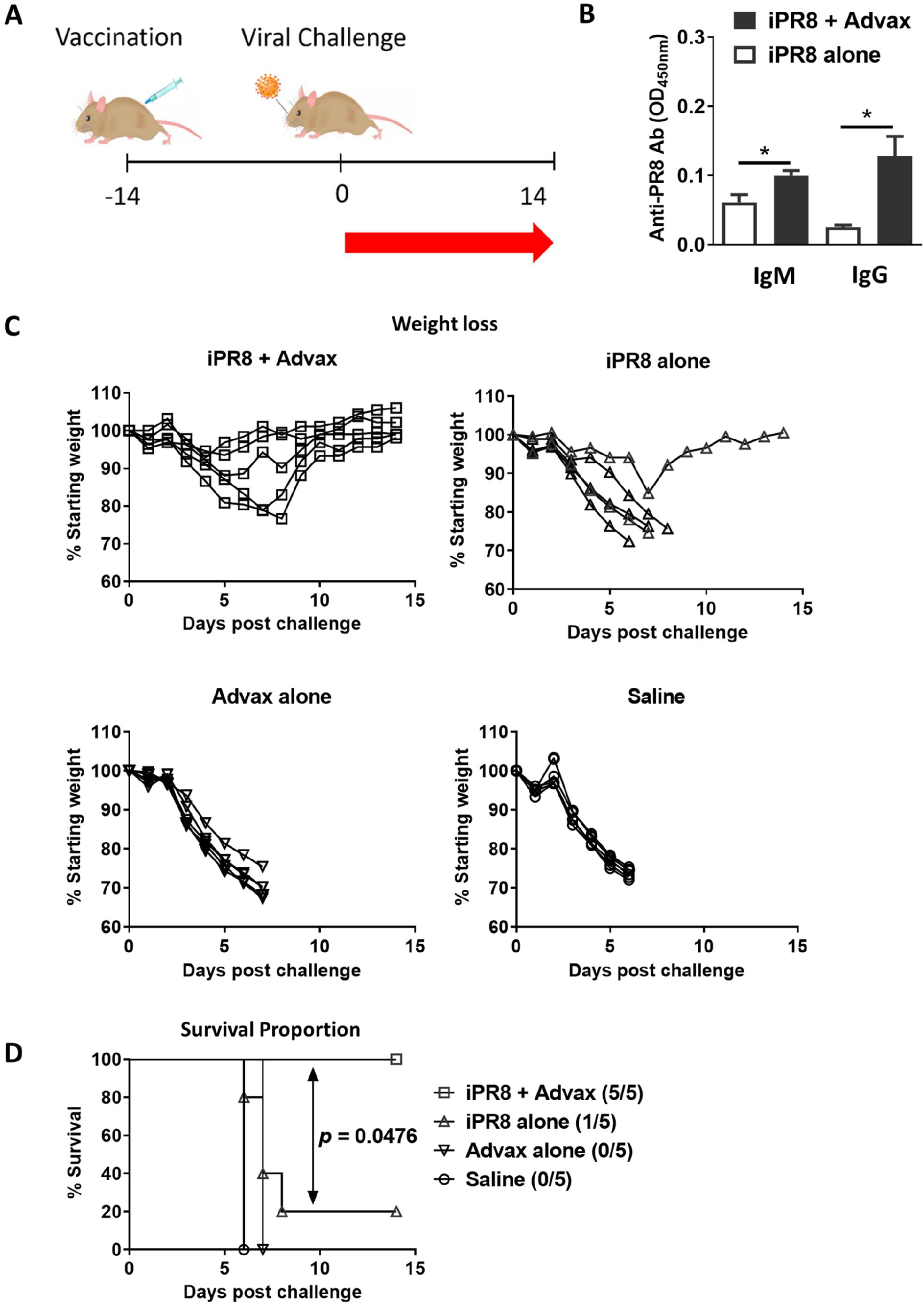
Effect of Advax adjuvant on immunogenicity and protective efficacy of a single-dose influenza vaccine injected 2 weeks before virus challenge. Female BALB/c mice were immunized i.m. with 300ng of inactivated PR8 (iPR8) influenza alone or together with 1 mg Advax adjuvant. Control group received Advax or saline alone. Blood samples were collected 2 weeks after immunization (B). All the mice were given an intranasal (i.n.) challenge with 8 x LD_50_ (560 PFU) PR8 virus. Shown are study design (A), anti-PR8 antibodies measured by ELISA (mean + SEM) (B), post-challenge bodyweight changes (mean ± SEM) (C), survival rate with number of survivors / number of challenged mice shown in parenthesis (D). Statistical analysis: Mann-Whitney test for ELISA data and log-rank (Mantel-Cox) test for survival rate. *, *p* < 0.05.

### Influenza vaccine formulated with Advax adjuvant formulation provides immune protection to infection immediately following vaccination

To evaluate whether the Advax-adjuvanted vaccine provides earlier immune protection than 2 weeks, immunised mice were challenged at progressively shorter times after immunisation, starting at 7 days, then moving to 3 days post-immunisation with complete protection continuing to be seen (Data not shown or supplementary data). We therefore asked whether there would be any protection if mice were challenged on the same day as immunisation. To ensure the results were applicable across different strains, both BALB/c and BL6 mice were challenged intranasally with 8 x LD_50_ of live PR8 virus immediately after intramuscular vaccination **(Figure 2A)**. The Advax-adjuvanted vaccine provided significant protection in both BALB/C and BL6 mice as measured by survival and post-challenge weight loss when compared to vaccine alone, with complete lethality in all control mice including those that received vaccine or Advax-adjuvant alone **(Figure 2B-C)**. The survival rate for same day administered Advax-adjuvanted vaccine was slightly higher in BALB/c mice (78.5%) than BL6 mice (66.6%).

**Figure 2.**
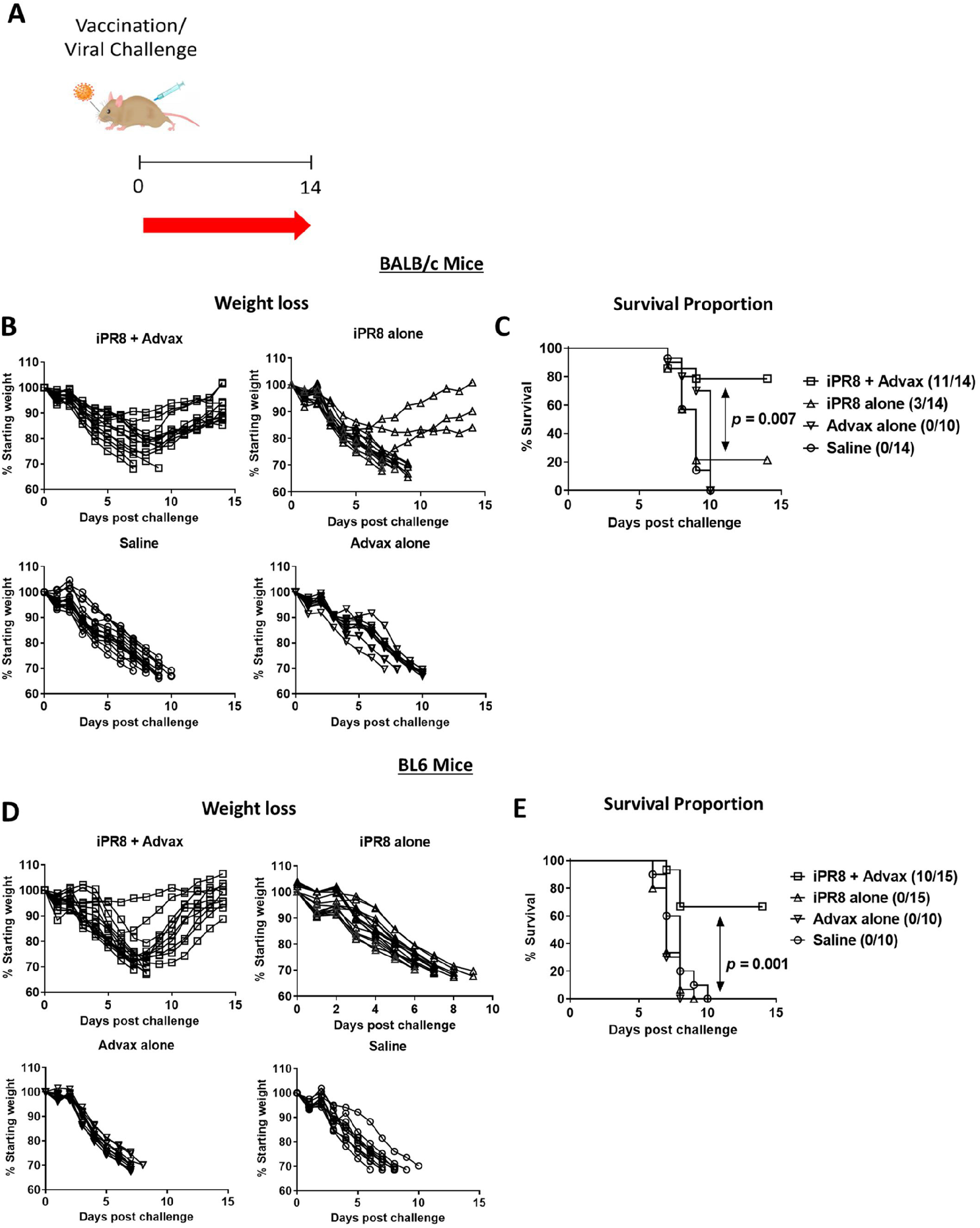
Effect of Advax adjuvant on immunogenicity and protective efficacy of a single dose influenza vaccine injected on the same day as viral challenge. Female BALB/c and BL6 mice were immunized i.m. with 5μg of inactivated PR8 (iPR8) influenza virus alone or together with 1mg Advax adjuvant. Control group received Advax or saline alone. All the mice were given an intranasal (i.n.) challenge with 8 x LD50 (560 PFU) PR8 virus. Shown is the study design (A), post-challenge bodyweight changes (mean ± SEM) (B, D), survival rate with number of survivors / numbers of challenged mice shown in parenthesis (C, E). Statistical analysis: MannWhitney test for ELISA data and log-rank (Mantel-Cox) test for survival rate. *, p < 0.05.

Next, we sought to identify the potential mechanism for the observed same day protection. Histopathology of lung sections at 5-7 days post challenge (dpc) showed significantly greater interstitial immune infiltration with peri-vascular cuffing and extensive alveolar thickening in animals receiving vaccine alone compared to Advax-adjuvanted vaccine **(Figure 3A)**. The lungs at 5-7dpc showed a progressive decrease in lung virus load in both vaccine groups, with a trend to a lower virus load in the Advax-adjuvanted group, although this difference did not reach statistical significance.

**Figure 3.**
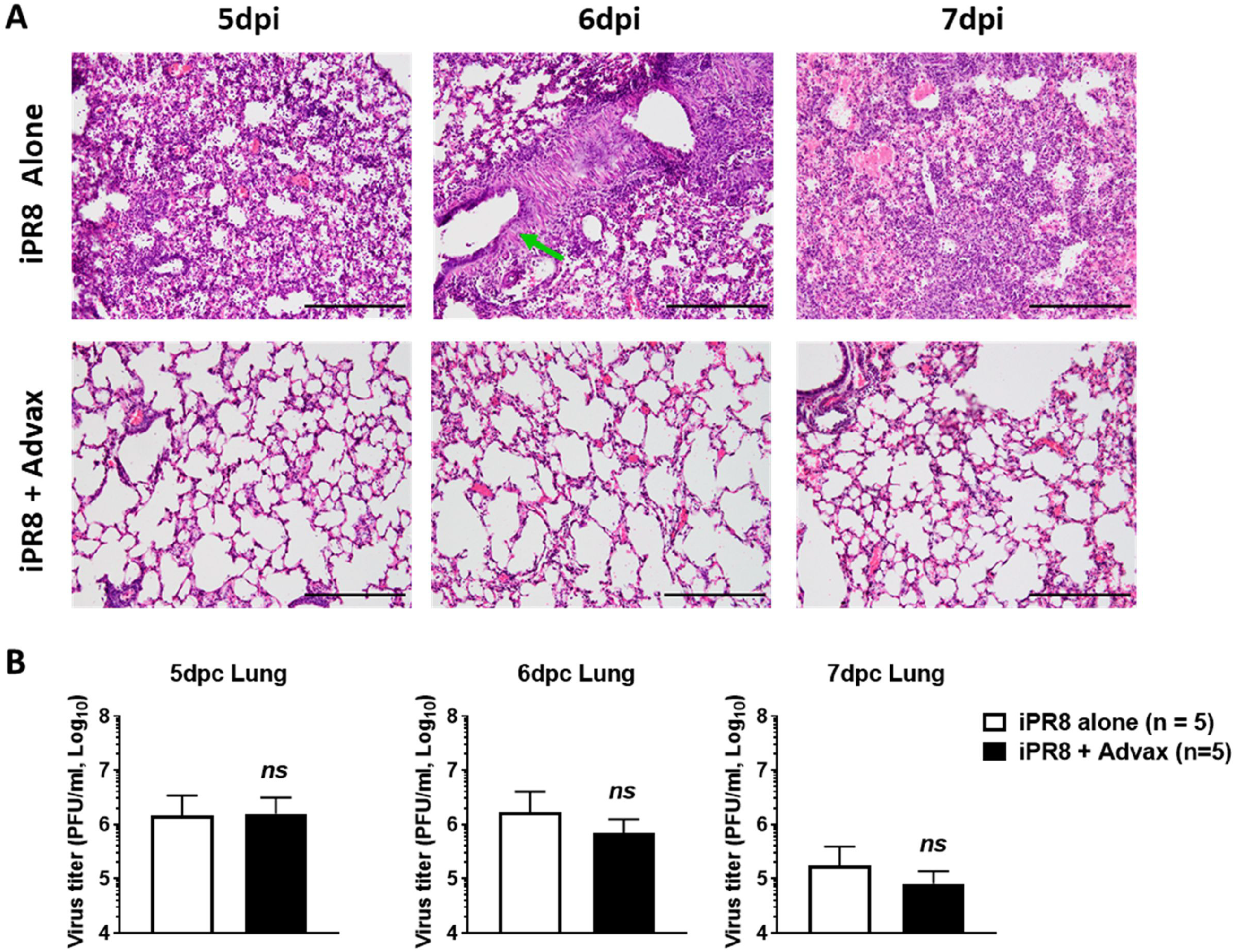
Lung pathology and virus titers in challenged mice. Female BALB/c mice (n = 15 / group) were immunized i.m. with 5μg inactivated PR8 (iPR8) virus with or without Advax adjuvant. Immediately after immunization, all the mice were challenged with 8 x LD50 dose of PR8 virus i.n. Five mice per group were euthanized daily between 5 to 7 days post-challenge (dpc) and blood and lung samples were collected. Lungs were fixed, sectioned, and stained with hematoxylin and eosin (H&E staining) and representative images from lung tissues of immunized and challenged mice are shown (A). The tissue sections were observed for pathological damages under light microscopy (10× Original magnification). Viral loads in the lung were determined in 20% homogenates by plaque assay (B). Each bar represents the mean ± SD (5 mice lungs per time point group). Statistical analysis: Mann-Whitney test. *; p < 0.05, ns; no significance.

**Figure 4.**
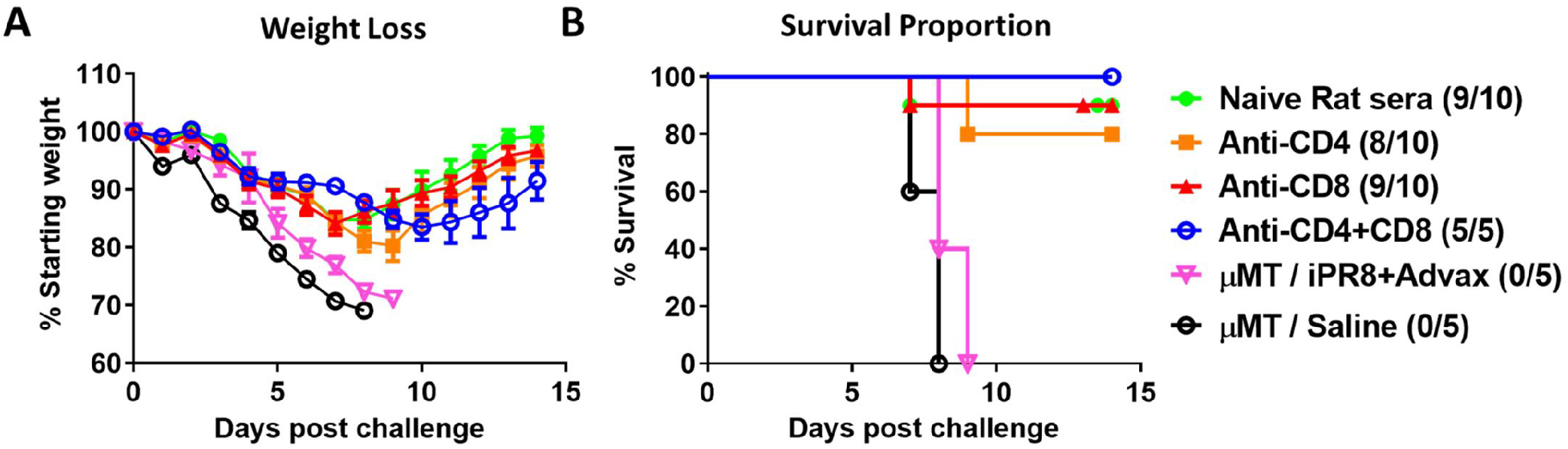
B-cells but not T-cells are essential for accelerated vaccine protection. Female B-cell-deficient (μMT) mice were immunized i.m. with 5μg inactivated PR8 (iPR8) virus with Advax. Female wild-type (WT) mice were treated with indicated T-cell depleting rat monoclonal antibodies or naïve rat sera i.p. on Day −3, −1 and +2 to immunization. All the mice were challenged with PR8 virus immediately after immunization. Shown are post-challenge bodyweight changes (mean ± SEM) (A) and survival rate with number of survivors / number of challenged mice shown in parenthesis (B).

### Accelerated Protection with Advax-adjuvanted vaccine is B-cell and antibody dependent

To define the mechanism responsible for the accelerated protection, B-cell deficient mice (μMT strain) and mice depleted of CD4+ and/or CD8+ T-cells using monoclonal antibodies were challenged with a lethal dose of PR8 virus immediately after immunization. The accelerated protection induced by Advax-adjuvanted iPR8 was completely abrogated in the μMT but not the T cell depleted mice, indicating the accelerated protection was B-cell and not T-cell dependent. Furthermore, passive transfer into naïve mice of sera collected 6dpc from immunised/challenged mice, also conferred protection **(Figure 5)**. These results indicate that the accelerated protection in Advax-adjuvanted iPR8 immunised mice is mediated by antigen-specific antibodies.

**Figure 5.**
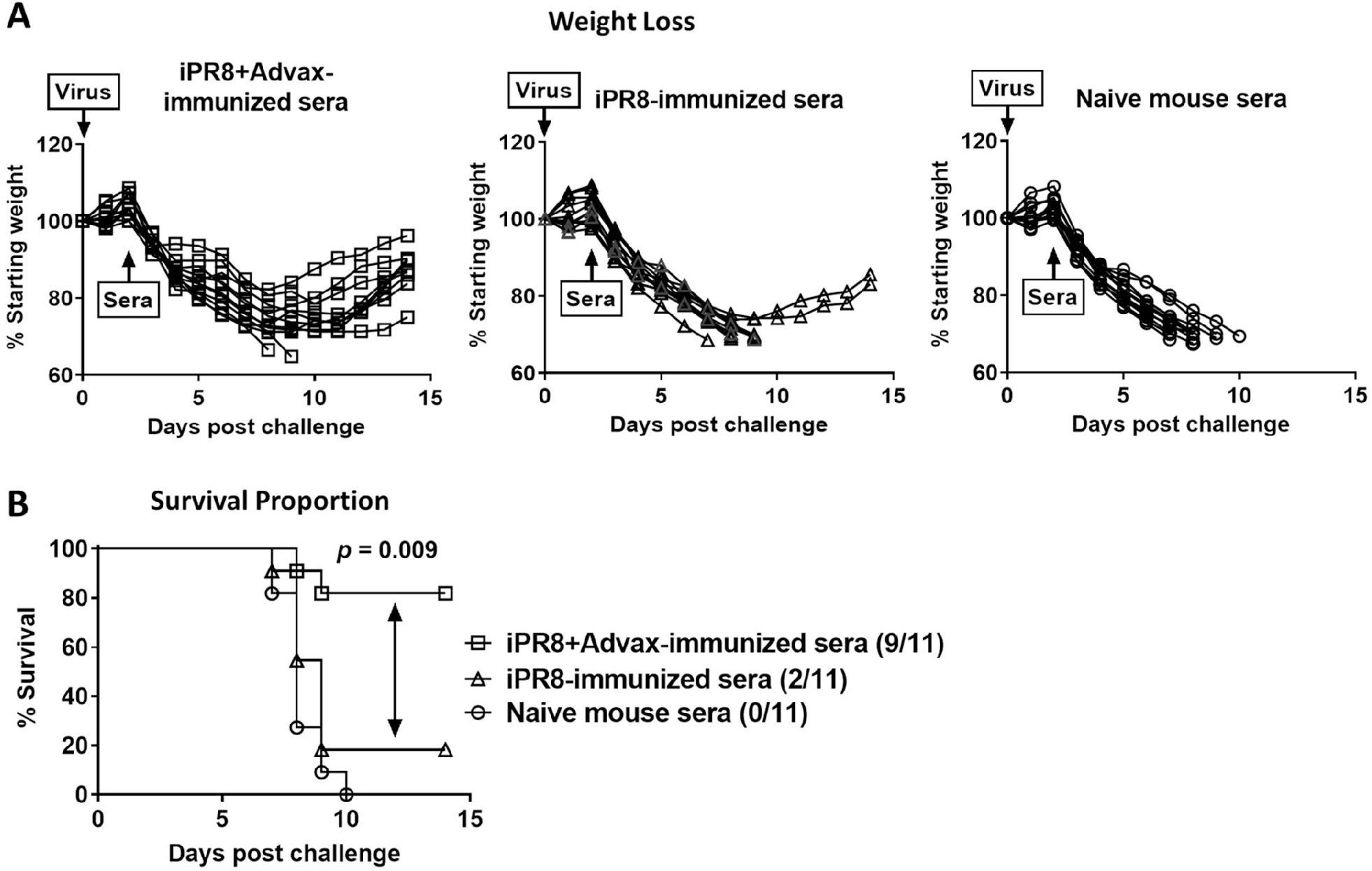
Passive transfer of day 6 immune sera confers protection to naïve mice. Sera for transfer were collected 6dpc from iPR8 alone-immunized & challenged or iPR8+Advax-immunized & challenged mice. Naïve BALB/c mice were intranasally infected with 8 x LD_50_ dose of PR8 virus and heat-inactivated sera were injected i.p. at 2dpc. Shown are post-challenge bodyweight changes (mean ± SD) (A) and survival rate with number of survivors / number of challenged mice shown in parenthesis (B). Combined results of 3 independent experiments.

### Advax adjuvant induces influenza-specific IgM that protects against viral challenge

Analysis of sera at 6dpc showed that influenza-specific IgM levels were significantly increased in the Advax-adjuvanted vaccine group compared to vaccine alone, while IgG levels were low with no major differences between groups. To confirm the specific isotypes involved in accelerated protection, size-exclusion chromatography was used to separate and pool the two main types of antibodies, IgG and IgM, from the 6dpc immunoglobulin fractions of mice simultaneously immunized and challenged **(Figure 6B-C)**. An HAI assay confirmed that 6dpc IgM was far more potent at neutralising influenza virus (1:640) when compared to 6dpc IgG (1:10) **(Figure 6D).** Finally, to confirm the immune IgM was responsible for accelerated protection in the Advax-adjuvanted iPR8 immunised mice, the purified 6dpc IgM was mixed with a lethal dose of PR8 virus used to challenge naïve mice. Whereas purified IgM from naïve mice provided no protection, 6dpc IgM from Advax-adjuvanted iPR8 immunised mice provided complete protection **(Figure 6E-F)**, thereby confirming the critical role of influenza-specific IgM in the accelerated vaccine protection.

**Figure 6.**
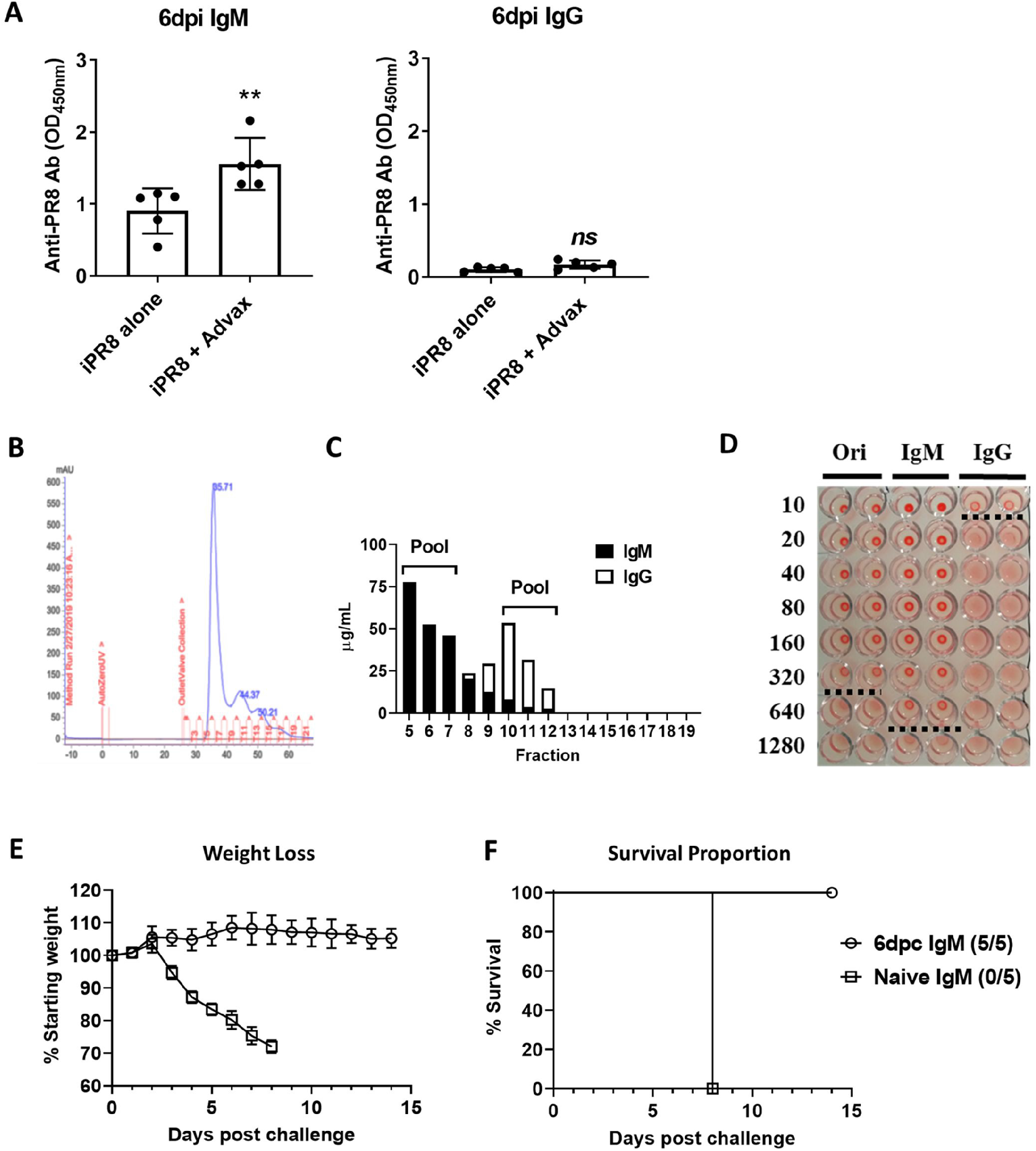
Day 6 immune IgM confers protection against influenza infection. Antigen-specific IgM and IgG antibody levels were determined by ELISA at 6pc (A). Immunoglobulin fraction obtained from iPR8+Advax-immunized mice at 6dpc was separated by size-exclusion chromatography to separate IgM and IgG (B). IgM and IgG concentrations were determined by sandwich ELISA (C) and protective antibody titers were determined by HI assay (D). Ori: original sera before column chromatography. IgM: Pooled fractions (Fr. 5 to 7 in 5B). IgG: Pooled fractions (Fr. 10 to 12 in 5B). Dotted lines show HI titer. Body weight (E) and survival rates (F) changes of mice received 1μg IgM mixed with lethal dose of PR8 virus.

## DISCUSSION

Over the last century, multiple influenza pandemics occurring every 20-30 years have resulted in a major loss of human life (21). The risk of an pandemics has increased in recent years due to expanding globalization, the rise in geographic disparate strains of virus and the existence of multiple animal reservoirs (domestic and wild) in which the pathogen can develop and undergo transformation (22). A vital tool in preparing for the next pandemic is the development of effective and fast acting vaccines. Traditional influenza vaccines generally have poor immunogenicity in the absence of an adjuvant (23). Advax is a novel polysaccharide-based adjuvant that is engineered from delta inulin into microparticles that has been shown to improve the immunogenicity and protection of influenza vaccines in animals and humans (9, 17). Immune protection can often take more than a month to develop and during this time, newly vaccinated individuals remain vulnerable to infection.

This study showed the ability using an appropriate vaccine strategy to accelerate the development of protective adaptive immunity against an otherwise lethal influenza infection. Notably, this protection was not mediated through innate immunity which is typically thought to be the only arm of the immune system able to respond this rapidly, as highlighted by the fact that the vaccine or Advax adjuvant on their own had no protective effect. In particular, it was the ability of the Advax adjuvant to induce rapid production of influenza-specific IgM with influenza neutralisation capacity that explained how the vaccine was able to outrace the virus and protect against an otherwise highly lethal infection. Lung pathology at 5-7 dpc in the protected animals group showed absence of pneumonia, contrasting with the high cell infiltration and thickening of the alveolar walls in the animals that just received the standard influenza vaccine without adjuvant. Notably, B-cells but not T-cells were essential for the accelerated protection.

This study reconfirms the oft forgotten vital role of IgM in early viral protection. IgM is the oldest antibody isotype at an evolutionary level and is the first antibody to be produced in an immune response to a foreign pathogen, with IgG appearing later following antibody class switching and affinity maturation (24). IgM possesses a number of attributes that make it a highly effective anti-viral defence system. It is highly multivalent with up to 10 Fab antigen binding sites enabling tight binding despite relatively low avidity of each individual Fab-antigen interaction. It is able to cross-link multiple viruses together and agglutinate them, thereby further impeding their ability to fuse with cells, being 100 to 10,000 times more effective than IgG in mediating agglutination (25). It binds and activates complement, and by binding the unique Fc receptor (FcμR) constitutively expressed on human lymphocyte cells can mediate effector function (26, 27). Finally, passively administered IgM have been shown to play an almost adjuvant-like role and enhance subsequent antibody responses to antigen challenge, whereas IgG antibodies suppress this response (28).

The importance of IgM antibodies in defence against influenza was highlighted by a study in which mutant mice unable to secrete IgM but normally expressing other Ig isotypes failed to control infection with influenza A/Mem71 (H3N1) (29). A recent study also found that IgM had higher potency and broader activity against influenza strains compared to IgG (16). Previous human influenza vaccine trials showed that Advax adjuvant enhanced influenza-specific IgM as well as IgG responses (19). Hence the importance of an adjuvant being able to rapidly induce antigen-specific IgM responses in order to provide accelerated protection is likely to have been overlooked in the past, given the almost exclusive emphasis in the past on vaccine-induced IgG. It is not clear whether other adjuvants might similarly be able to able to enhance early IgM responses or whether this could be an exclusive property of delta inulin, a unique adjuvant that has anti-rather than pro-inflammatory actions thereby distinguishing it from more traditional adjuvants.

A limitation of this study is that there can be differences in immune responses of mice and humans (30) so similar findings in additional species, such ferrets or non-human primates, would strengthen the study findings. Ferrets and non-human primates in addition to more closely matching humans in terms of physiology and disease pathogenesis also have increased genetic diversity (31). In addition to extending the findings to additional species, planned future studies will also examine whether this effect of Advax adjuvant on accelerating vaccine protection extends to additional viral pathogens, including SARS-CoV-2. These studies will also examine whether the persistence of such antigen-specific IgM antibodies and whether a similar accelerated IgM immune response is observed in the event of rechallenge with the virus at a later timepoint. The results nonetheless provide proof of concept that the inclusion of Advax adjuvant to inactivated influenza vaccine can help provide protection from concomitantly acquired infection.

In summary, our study highlights the importance of IgM to early virus protection, demonstrating that with the use of an appropriate adjuvant such as Advax it is possible to dramatically accelerate the production of antigen-specific IgM so as to enable vaccine protection against a co-administered lethal virus challenge. These findings could have important ramifications for pandemic vaccine design, and reinforce the importance of research into novel adjuvants, plus the regulation and role of IgM in viral defence.

## MATERIALS AND METHODS

### Experimental Animals

Female BALB/c, C57BL/6 (BL6) and B cell-deficient μMT mice (6 to 10 weeks old) were supplied by the College of Medicine and Public Health Animal Facility of Flinders University of South Australia. Virus challenge experiments were performed in a PC2 animal laboratory. Mice were immunized by a single intramuscular (i.m.) injection in the hind thigh with formalin-inactivated influenza A/Puerto Rico/08/34 virus (iPR8) (Charles River, Norwich, CT, U.S.A.) with or without 1mg Advax adjuvant. Advax is made from microparticulate delta inulin and was obtained from Vaxine Pty Ltd. (Adelaide, SA, Australia). The study was carried out in strict accordance with the Australian Code of Practice for the Care and Use of Animals for Scientific Purposes (2013). The protocol was approved by the Animal Welfare Committee of Flinders University.

### Cell Lines

MDCK cell line was obtained from the American Type Culture Collection (ATCC, CCL-34) and maintained in DMEM supplemented with GlutaMAX, 100U/ml of penicillin, 100μg/ml of streptomycin and 5% fetal calf serum (FCS; HyClone, GE Healthcare, South Logan, UT, U.S.A.) at 37°C in a humidified atmosphere with 5% CO_2_.

### PR8 Challenge of mice

For influenza challenge studies, unimmunized control groups were routinely included to ensure that the H1N1/PR8 challenge dose was completely lethal. Mice were infected by intranasal (i.n.) administration of 50μl of saline containing virus under ketamine (75mg/kg) and medetomidine (1mg/kg) anesthesia. The virus challenge dose used for wild-type (WT) mice was 8 x LD_50_ whereas a lower 1 x LD_50_ dose was used for μMT mice because of their higher susceptibility to virus lethality when compared to WT animals. One LD_50_ of PR8 virus was calculated as 70 PFU in adult BALB/c mice. Mice were evaluated daily and scored for individual symptoms, as described elsewhere (7). Mice were euthanized if they developed a clinical score of 6 or if they lost in excess of 30% of their pre-challenge body weight.

### Sample collection

Blood samples were collected from facial vein or cardiac puncture under anesthesia for serum transfer studies. Serum was separated by centrifugation and was stored at −20°C prior to use. Five mice from each group were sacrificed by cervical dislocation for histology and titration of residual viruses in the lungs. Lungs were aseptically removed from mice after euthanasia. Left lungs were fixed in neutral buffered formalin (10%) for 24–48 h. Right lungs were homogenized in Dulbecco’s Modified Eagle Medium (DMEM; Thermo Fisher Scientific, Scoresby, VIC, Australia) by a Mini-Beadbeater-16 homogenizer (Bio Spec Products Inc., Bartlesville, OK, U.S.A.). Final volume of lung homogenates was adjusted by DMEM to obtain 20% (wt/vol) homogenate and clarified by centrifugation at 2000 x g for 10 min. Supernatants of homogenized lungs were aliquoted and stored at −80°C until analyzed.

### Virus titration

MDCK cells were plated in 6-well plates (Greiner Bio-One) and incubated overnight at 37°C with 5% CO_2_. Serial 10-fold dilutions of the samples were made in DMEM containing 3% FCS. The plates were washed once with PBS and the 400μl of inoculum was added to each well. Following a 1 h adsorption at RT, the viral inoculum was removed from the cell monolayer. The cells were washed once with PBS and then overlaid with 2ml/well of 1.2% Avicel RC-591 (FMC Biopolymer, Philadelphia, PA, U.S.A.) in MEM medium containing 2μg/ml TPCK-treated trypsin (Sigma-Aldrich) and 3% FCS. After 3 days of incubation at 37°C, the overlay was removed and washed with PBS. The cells were fixed with 50% (v/v) Aceton/Methanol for 10 min at RT. The cells were stained with an anti-Influenza NP mouse monoclonal antibody (BEI NR-4544) and biotinylated anti-mouse IgG1 (Abcam, Cat. No. ab 97238) mixed with HRP-conjugated Streptavidin (BD, Cat. No. 554066). The plaques were visualized with SIGMAFAST DAB with Metal Enhancer (Sigma-Aldrich).

### Histological examination

Lungs were fixed in 10% formalin in PBS and embedded in paraffin wax and sections were cut at 10μm thick. Sections were then stained with hematoxylin and eosin (H&E). Slides were scanned by light microscopy.

### Antibody assays

Influenza-specific antibodies and mouse IgM and IgG concentrations were determined by ELISA. iPR8 antigen (1μg/ml) was used to coat each well of Microlon medium-binding 96-well ELISA plates (Greiner Bio-One, Cat. No. 655001). For quantification of mouse IgM and IgG, plates were coated with goat anti-mouse IgM (Sigma-Aldrich, Cat. No. M8644) or goat anti-mouse IgG (Sigma-Aldrich, Cat. No. M1397) at 1μg/ml. Plates were blocked by 200μl/well of blocking buffer (0.2% casein in 50mM Tris-HCl, pH8.0 and 0.05% Tween 20) for 1 h at RT. After removing blocking buffer, 100μl of samples in blocking buffer were added to wells and incubated for 2 h at RT. Purified mouse IgM and IgG1 (BD Pharmingen, Cat. No. 557275 and 554121) were used as standard. Biotinylated anti-mouse IgM (Abcam, Cat. No. ab97228) and IgG (Sigma-Aldrich, Cat. No. B7022) were used as secondary antibodies. After 6 wash with PBS/Tween 20 (0.05%), 100μl of diluted secondary antibody was added to each well with 1:2000 dilution of HRP-conjugated Streptavidin (BD) and incubated for 1 h at RT. After another 6 wash with PBS/Tween, plates were incubated with 100μl of freshly prepared TMB substrate (KPL, SeraCare, Milford, MA, U.S.A.) for 10 min. Reaction was stopped by adding 100μl of 1M Phosphoric Acid. The optical density was measured at 450nm (OD_450nm_) using an automated spectrophotometer plate reader (VersaMax, Molecular Devices, CA, U.S.A.) and analyzed using SoftMax Pro Software. Hemagglutination inhibition (HI) assay was performed as described previously using 1% guinea pig red blood cells as described (7).

### *In vivo* B-cell and T-cell depletion assay

For *in vivo* depletion, 200μg anti-CD4 (Clone GK1.5) and 250μg anti-CD8 (Clone 53-6.72) (both from BioXCell, West Lebanon, NH, U.S.A.) or 450μg naive rat IgG prepared from naive rat serum by 30% ammonium sulfate precipitation, were injected i.p. at days −3, −1 and +2 of immunization and virus challenge. Depletion was verified by flow cytometry using peripheral blood cells treated with RBC lysis buffer and rat anti-mouse CD16/CD32 (Clone 2.4G2) then stained with APC-anti-mouse CD4 (Clone RM4-5) and PE-anti-mouse CD8a (Clone 53-6.72) (all from BD).

### IgM and IgG separation

Immunoglobulins were precipitated from day 6 challenge sera or naïve mouse sera by 50% ammonium sulfate precipitation and then IgM and IgG were separated by HiPrep Sephacryl S-200 H chromatography columns (GE Healthcare Australia, Silverwater, NSW, Australia).

### Immunoglobulin fractions *in vivo* protection assay

One microgram of IgM obtained from day 6 serum of mice immunized and challenged on the same day or from naive animals was mixed with the challenge dose of PR8 virus and incubated at room temperature for 30 min. Naive mice were then inoculated i.n. with a mixture of the day 6 IgM and virus and then monitored for their bodyweight and sickness scores till day 14 post virus challenge.

### Statistical analysis

Antibody titers, body weight changes and residual virus titers in the lungs were compared among different groups of mice by Mann-Whitney test using GraphPad Prism version 5.02 (GraphPad Software, San Diego CA, U.S.A.). For animal survival data, survival curves were created using the Kaplan-Meier method and statistical analyses of survival rate was performed with two-sided Fisher’s exact test. For all comparison, *p* < 0.05 was considered to represent a significant difference.

## Acknowledgments

We thank Chun Hao Ong, Mariah Castignani and Kodie Noy for assistance with animal husbandry and preparation of lung sections. The following reagent was obtained though BEI Resources, NIAID, NIH: Monoclonal Anti-Influenza A Virus NP, Clone IC5-1B7 (produced in vitro), NR-4544. This work was supported by funding from National Institute of Allergy and Infectious Diseases of the National Institutes of Health under Contracts. HHS-N272201400053C, HHS-N272200800039C and U01-AI061142.

## Authors Contributions

YHO and NP designed the research; YHO and LL performed the research; YHO, JB and NP analyzed data; YHO, LL, JB and NP wrote the paper. All authors have read and approved the final manuscript.

---

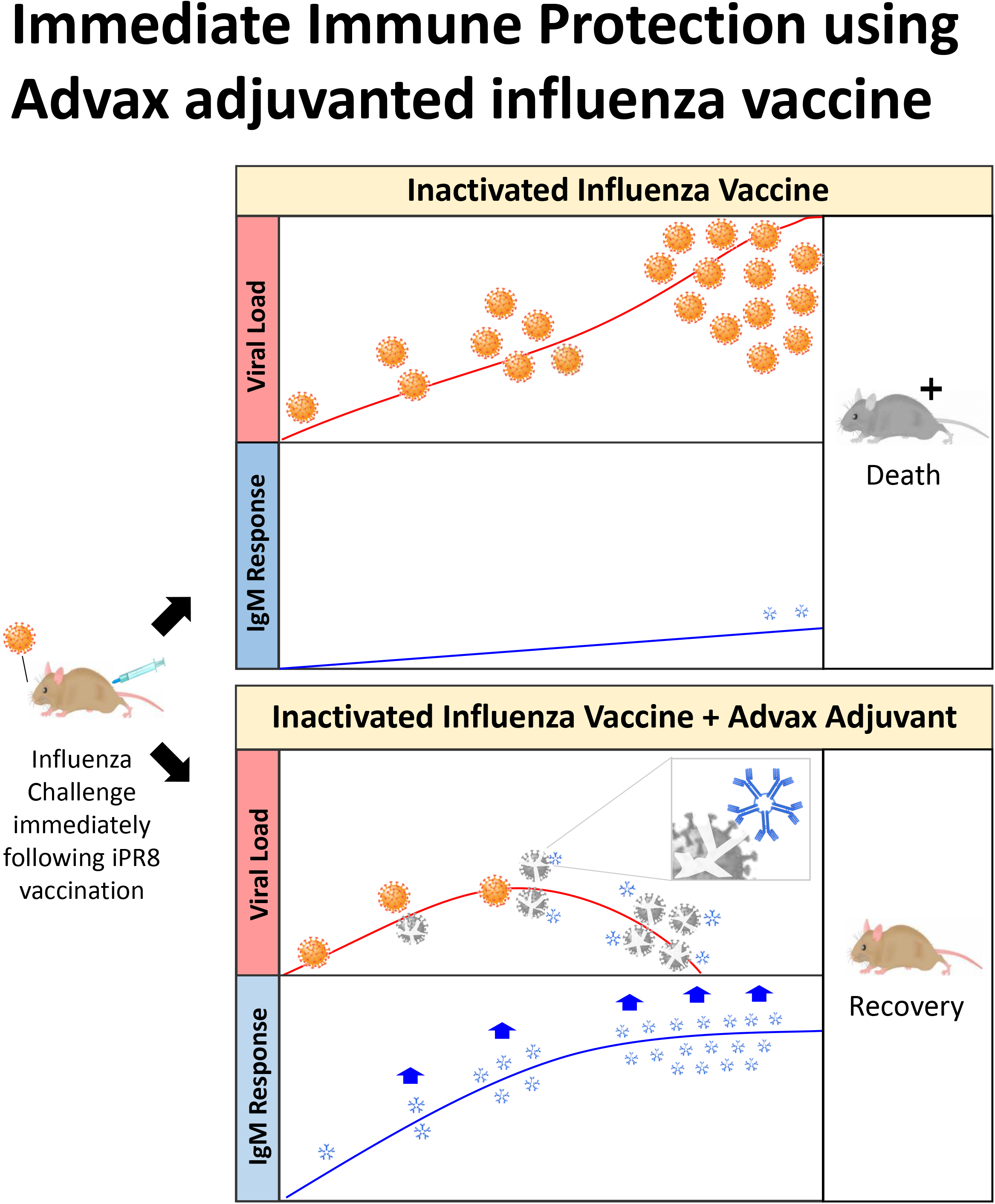

